# COSSMO: Predicting Competitive Alternative Splice Site Selection using Deep Learning

**DOI:** 10.1101/255257

**Authors:** Hannes Bretschneider, Shreshth Gandhi, Amit G Deshwar, Khalid Zuberi, Brendan J Frey

## Abstract

**Motivation:** Alternative splice site selection is inherently competitive and the probability of a given splice site to be used also depends strongly on the strength of neighboring sites. Here we present a new model named Competitive Splice Site Model (COSSMO), which explicitly models these competitive effects and predict the PSI distribution over any number of putative splice sites. We model an alternative splicing event as the choice of a 3’ acceptor site conditional on a fixed upstream 5’ donor site, or the choice of a 5’ donor site conditional on a fixed 3’ acceptor site. We build four different architectures that use convolutional layers, communication layers, LSTMS, and residual networks, respectively, to learn relevant motifs from sequence alone. We also construct a new dataset from genome annotations and RNA-Seq read data that we use to train our model.

**Results:** COSSMO is able to predict the most frequently used splice site with an accuracy of 70% on unseen test data, and achieve an *R*^2^ of 60% in modeling the PSI distribution. We visualize the motifs that COSSMO learns from sequence and show that COSSMO recognizes the consensus splice site sequences as well as many known splicing factors with high specificity.

**Availability:** Our dataset is available from http://cossmo.deepgenomics.com.

**Supplementary information:** Supplementary data are available at *Bioinformatics* online.

## 1 Introduction

RNA splicing has long been known as a main driver of transcriptional diversity and a large number of regulatory mechanisms have been described. More recently, efforts have shifted from describing isolated regulatory mechanisms to building computational models, known as *splicing codes* (Wang and Burge, 2008), which can predict splice site usage from sequence directly or from a library of hand-engineered, sequence-derived features. Mis-splicing is also one of the leading mechanisms for genetic disease (Scotti and Swanson, 2015), which creates an important need for algorithms to accurately predict splicing *in-silico*, for example, to score the splicing effect of variants or design splicing-modulating therapies.

The first practical splicing code was introduced by Barash *et al*. (2010) and predicted tissue differences of cassette splicing events in mouse. Subsequent versions of the splicing code introduced a Bayesian neural network and predicted absolute splicing levels (Xiong *et al*., 2011). Since then, Bayesian neural networks (Xiong *et al*., 2015) and deep neural networks (Leung *et al*., 2014) have further improved on the state of art in predicting exon skipping.

These splicing codes have so far all modeled cassette splicing events. Busch and Hertel (2015) present a model that uses support vector machines to predict whether an exon is constitutively spliced, undergoes alternative 5’ or 3’ splice-site selection, or is an alternative cassette-type exon. However, their model does only predict the class of an exon, not the isoform or splice site utilization levels.

Splice sites are often seen as belonging to discrete categories such as constitutive and alternative sites. However, these are functional descriptors rather than properties of the splice site itself. For example, a constitutive splice site may be close to a cryptic splice site that is normally not recognized. However, a variant may activate the cryptic site such that the previously constitutive splice site is now alternatively used (Vaz-Drago *et al*., 2017). There are therefore two aspects that may affect the utilization of a splice site: The inherent strength of the splice site itself, determined by, for example, nearby binding motifs for splicing enhancers. But the utilization also depends on the strength of neighboring splice sites that could be used alternatively and compete with each other for recognition by the spliceosome. We therefore believe it is necessary to model the *competitive* aspect of splice site selection in addition to modeling the splice site itself.

Another development in the field is to learn features from sequence directly, rather than constructing feature libraries by hand. Constructing feature sets by hand can be extremely laborious and computational constraints may limit the size of the dataset that can be used for training. Convolutional networks that learn from sequence directly have already been used successfully in learning transcription factor binding motifs (Alipanahi *et al*., 2015), predicting the function of non-coding DNA (Quang and Xie, 2016), and predicting a large number of epigenetic and transcriptional profiles (Kelley *et al*., 2018).

In this paper, we present a new splicing code that we call COSSMO, for *competitive splice site model*. COSSMO is more general than previous splicing models and is capable of predicting a usage distribution of multiple splice sites, conditional on a constitutive opposite site. In particular, we can model the usage distribution of multiple alternative acceptor sites conditional on a constitutive donor site or vice versa.

We train two versions of this model, one for alternative acceptor sites and one for alternative donor sites. Figure 1a shows an example of an alternative acceptor event. In the case of alternative acceptors we always condition on a constitutive donor site. In this example we model four alternative acceptor sites, but the model can dynamically adapt to any number of sites. COSSMO uses these alternative sites along with the constitutive site as input and predict a discrete probability distribution over the alternative sites indicating the frequency with which they are selected.

Figure 1b shows an analogous case of alternative donor selection, where multiple donor sites are selected amongst to be spliced to a constitutive acceptor site.

**Fig. 1.**
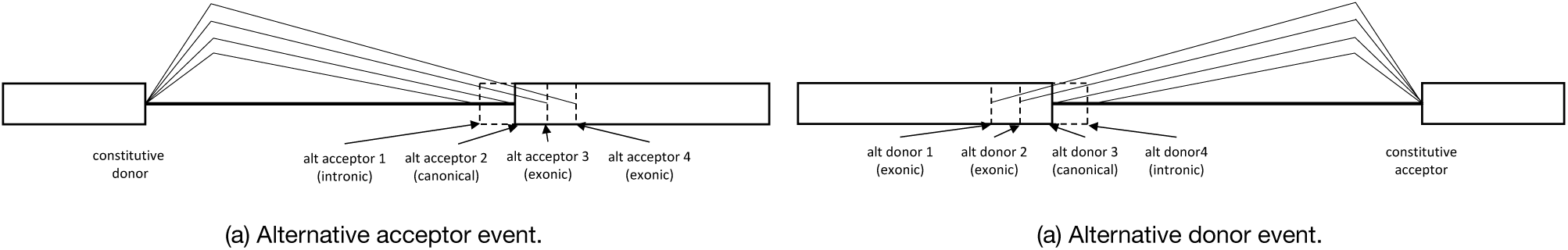
Illustration of alternative acceptor and donor events modeled by COSSMO.

## 2 Dataset Construction

In this section we describe how to construct a genome-wide dataset of quantified *percent selected index* (PSI) values of splicing events from genome annotations and RNA-Seq data from the *Genotype-Tissue Expression Project* (GTEx) (GTEx Consortium, 2013). An *event*, in this context, means a training example consisting of multiple putative splice sites that can be used alternatively. The PSI value is the frequency with which each splice-site is selected versus all other splice sites in the same event with PSI values for each event summing up to one.

We create two datasets: one of alternative acceptor events conditional on constitutive donor sites and one of alternative donor sites conditional on constitutive acceptor sites.

We mine an initial set of splicing events from genome annotations and then expand it with *de-novo* splicing events detected from the aligned GTEx RNA-Seq data. We further augment this set with non-splice sites. This is to increase the variability of features seen by the model and to help train the model to discriminate between splice sites and non-splice sites.

Each example in the alternative acceptor dataset consists of multiple putative acceptor splice sites and a constitutive donor splice site. In the alternative donor dataset, each example consists of multiple putative donor splice sites and a constitutive acceptor splice site.

Following the construction of these events, we quantify the PSI distribution for each example from RNA-Seq junction reads using the Bayesian bootstrap estimation method from Xiong *et al*. (2016). This allows us to quantify the uncertainty of our PSI values by estimating a posterior distribution.

### 2.1 Genome Annotations

We use the Gencode v19 annotations (Harrow *et al*., 2012) to construct our initial dataset. We start with a dataset of all exon-exon junctions by creating a training example for each annotated donor site and adding all acceptor sites that splice to this donor to the training example as alternative acceptors of type *annotated*.

Then we construct the initial alternative donor dataset analogously by finding the set of donors that splice to each acceptor site.

Figure 2 shows histograms of the number of alternative acceptors per donor and alternative donors per acceptor, respectively. As can be seen from the histogram, we do not only include splice sites that are annotated as alternatively spliced, but also constitutive exons. We do this to learn a general model of splice site strength. Summary statistics of the dataset are given in Table 1.

**Table 1.** Dataset statistics.

|  | Acceptor | Donor |
| --- | --- | --- |
| Number of events | 173,164 | 169,650 |
| Avg # of annotated sites/event | 1.158 | 1.184 |
| Avg # of *de-novo* sites/event | 3.744 | 3.696 |
| Avg # of decoy sites/event | 22.451 | 11.877 |
| Avg # of negative sites/event | 61.443 | 56.612 |

Each splicing event or training example consists of multiple putative splice sites each belonging to one of four types. *Annotated* sites are found in the Gencode v19 genome annotations. *De-novo* are not annotated but have RNA-Seq support in GTEx. *Decoy* sites have a MaxEntScan score greater than 3.0, but are not annotated and have no read support. *Negative* sites are random genomic locations.

**Fig. 2.**
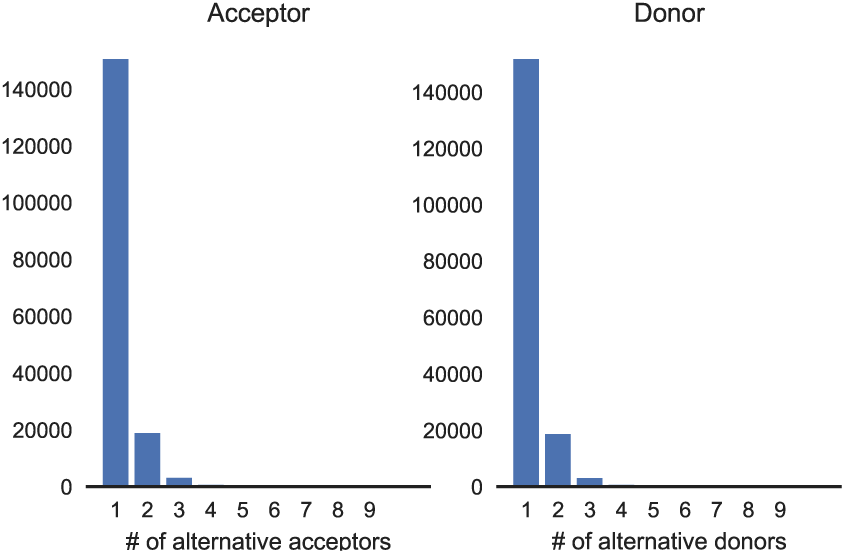
Histogram showing the complexity of annotated splicing events. For each annotated donor we count how many acceptors splice to it (left) and for each annotated acceptor we count how many donor sites are spliced to it (right).

**Fig. 3.**
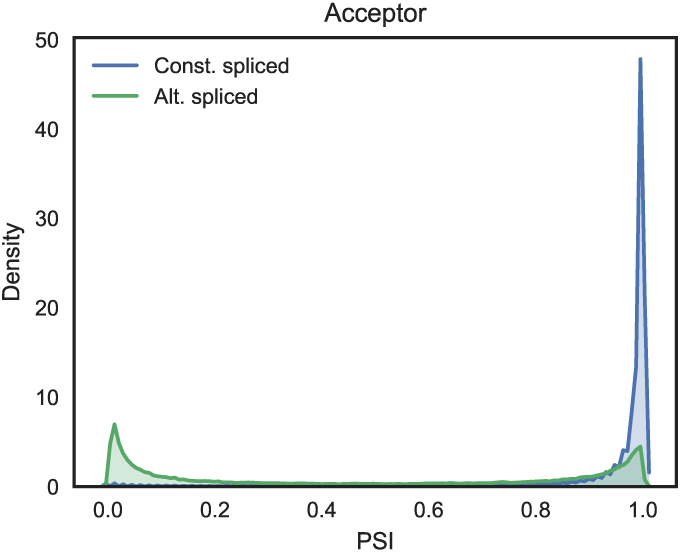
Kernel density estimator plot of PSI values in constitutively versus alternatively spliced events. Constitutively spliced events are defined as having exactly one annotated splice site while alternatively spliced events are all events with two or more annotated sites. Only acceptor dataset is shown.

### 2.2 *De-novo* Splice Sites from Gapped RNA-Seq Alignments

Genome annotations provide us with an initial set of alternative splicing events to form the core of our dataset. Next, we add *de-novo* splice sites from the GTEx RNA-Seq data. We start by aligning the GTEx RNA-Seq reads to the genome using the HISAT2 aligner (Kim *et al*., 2015). Next, we iterate through the dataset we built in Section 2.1. For the alternative acceptor dataset we iterate through all donor sites and for each site, we add all junction reads in GTEx that use this site as one end of a gapped alignment. We then add the other side of the gapped alignment as an alternative acceptor to the example. The procedure for adding *de-novo* splice sites to the alternative donor dataset is analogous: we iterate over all acceptor sites, find junction reads that use that acceptor site as one side of a gapped alignment and add the other end of the gap as an alternative donor site.

Some filtering of those *de-novo* sites is necessary both because the RNA splicing process itself is noisy and because sequencing and alignment introduce their own biases and artifacts, which would otherwise result in large numbers of low-certainty splice sites. We only utilize *de-novo* sites that are observed in at least two tissues from at least two subjects. This procedure results in a large expansion of possible splice sites, adding an average of 3.74 *de-novo* splice sites to each acceptor event, and 3.70 *de-novo* sites to each donor event (Table 1).

### 2.3 Negative Splice Site Examples

The above procedure gives us a dataset of high-confidence annotated and *de-novo* splice sites. However, we supplement this dataset with additional, verified, *non-splice sites*. This is done to increase the amount of variation in the features when training the model. For example, the core dinucleotides `GU` at the donor site and `A`G at the acceptor site are almost always a necessary feature for splicing to occur and will, therefore, be present in all examples of real splice sites (apart from examples using the *minor spliceosome*, Turunen *et al*. (2013)). If we trained a splicing model only on true splice sites, the model might learn to ignore the core dinucleotide motif since it will not provide any signal to the model. However, if we add some examples of non-splice sites to the dataset, then the model has an opportunity to learn that the presence of a core dinucleotide motif is necessary for recognition of the site. We call examples like this *negative sites* and we select them by sampling random locations from a region that (when constructing an alternative acceptor dataset) starts 20nt downstream of the donor site and ends 300nt downstream of the most distant alternative acceptor site.

Secondly, the genome also contains a large number of *decoy splice sites*. These are sites that look very similar to real splice sites and in many cases have a core dinucleotide motif, but are nevertheless not used as splice sites. For example, they may lack other necessary features such as a polypyrimidine tract or a branch point, or are adjacent to a silencer motif. These cases are also beneficial to include in our training set because they help the model detect the more subtle signals beyond the consensus sequence that are necessary for a splice site to be recognized by the spliceosome. We sample decoy splice sites by using MaxEntScan (Yeo and Burge, 2004) to scan the intron and exon for any sites that have a score greater than 3.0. We then remove from this set all sites that are either annotated as splice sites or if there are any junction reads aligned to them in the GTEx RNA-Seq data. The remaining sites are, therefore, locations that are assigned a high score by MaxEntScan and look very similar to true splice sites but for which there is no evidence that they are ever used as splice sites from annotations or GTEx.

On average, each acceptor event contains 22.45 decoy sites and 61.44 negative sites. Each donor event has, on average, 11.88 decoy sites and 56.61 negative sites (Table 1).

### 2.4 PSI Estimation

After building the datasets of alternative acceptors and donors, we quantify the frequency with which these sites are used. Our method for estimating PSI values is based on counting the number of junction reads that are aligned to a splice site pair.

Assuming we are interested in a constitutive donor site with multiple alternative acceptor sites, we could obtain a naive PSI estimate by counting the number of junction reads spanning from the donor to each alternative acceptor and normalizing those values to obtain a probability distribution. In practice, it is well known that RNA-Seq alignments exhibit many biases such as read stacks and other positional biases that make such naive estimates unreliable. Many methods have been developed to ameliorate these issues as well as to obtain confidence estimates of PSI values. We use the positional bootstrap method by Xiong *et al*. (2016). This method estimates a non-parametric posterior distribution of PSI using a Bayesian positional bootstrap procedure. For the purposes of this paper, we do not focus on tissue differences in splicing and instead investigate splicing regulation more generally. Therefore, to estimate PSI from the GTEx RNA-Seq data, we pool the reads from all GTEx samples to estimate an average PSI across all subjects and tissues, excluding cancer tissues and cell lines. This means for any biological replicates as well as different tissues we simply use all available reads at any given position for our PSI estimates.

## 3 Model

### 3.1 Motivation

The primary design goal of COSSMO is to construct a model that can dynamically predict the relative utilization of any number of competing putative splice sites. We achieve this by a network architecture that uses the same weights to score all splice sites and uses a *dynamic softmax* function at the output to adjust the size of the output layer to the number of alternative splice sites in the example.

For each putative splice site, the inputs to the model are DNA and RNA sequences from 80nt wide windows around the alternative splice sites and the paired constitutive splice site, as well as the intron length (distance between acceptor and donor sites). The model’s output is a PSI estimate for each putative splice site.

We first present the *acceptor model*, which predicts the PSI of multiple acceptor sites conditional on a constitutive donor site, but the donor model is constructed completely analogously by swapping *acceptor* and *donor*, as well as *3’* and *5’* everywhere.

Given a constitutive donor splice site *const* and *K* alternative acceptor splice sites *alt*_1_, *alt*_2_, …, *alt_K_*, COSSMO is a function *f* that predicts the probability of selecting the *k*-th splice site conditional on the donor site,

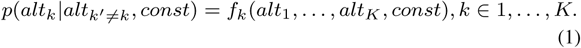

### 3.2 Features

All sequence features are represented as *one-hot* encoding using four channels that represent the four possible nucleotides.

For each splice site we extract the following sequence inputs:

> *Alternative DNA sequence:* This is the pre-splicing sequence from a 80nt window around the alternative acceptor site.
>
> *Constitutive DNA sequence:* This is the pre-splicing sequence from a 80nt window around the constitutive donor site (and thus it is the same for all splice sites in the same event).
>
> *mRNA sequence:* This is the spliced mRNA sequence obtained by concatenating 40bp of exonic sequence upstream of the donor site and 40nt of exonic sequence downstream from the acceptor site.

To these sequence inputs we add a single feature representing the intron length, obtained by computing the distance between the constitutive donor and alternative acceptor site and normalizing this distance by the mean and standard deviation of the length of an intron in the human genome (Hong *et al*., 2006) to make it more numerically stable.

COSSMO consists of two primary components: a *scoring network* that produces a scalar unnormalized score for a single splice site and a softmax layer that normalizes scores from multiple scoring networks.

The requirements for the scoring network are that it takes a single splice site’s sequence as input and produces a single scalar score. The output layer simply accepts the scalar scores as input, normalizes them, and then outputs the predicted PSI distribution over the alternative splice sites. The high-level architecture is illustrated in Figure 4.

**Fig. 4.**
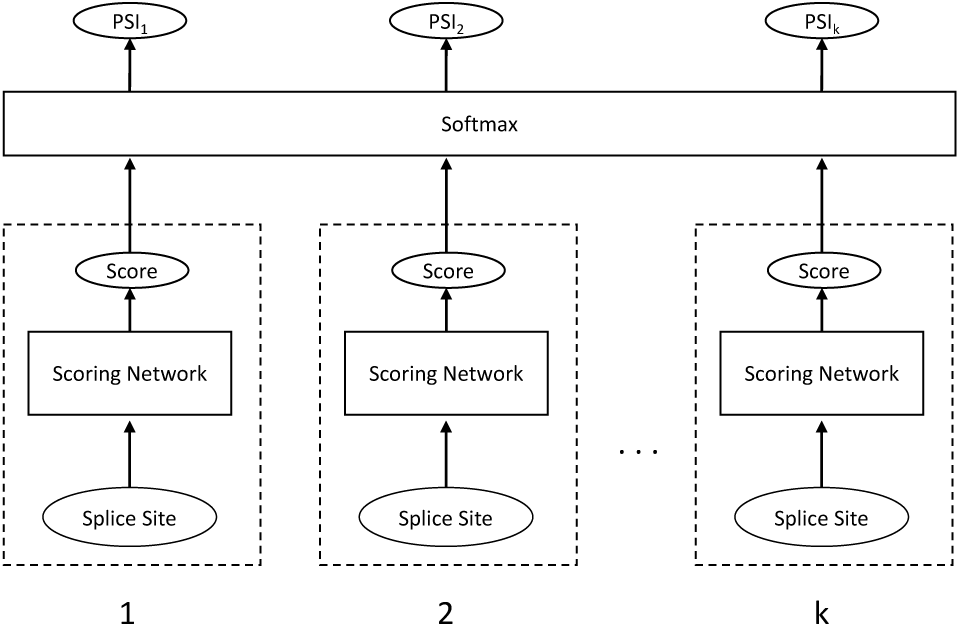
COSSMO Architecture. A score is computed for each putative splice site separately using identical sub-networks with weight sharing. The scores are then normalized with a softmax layer, allowing the number of splice sites to vary.

While the scoring network’s task is to predict a scalar score for each alternative splice site, we also need an output layer that normalizes those scores to obtain a valid probability distribution over the splice sites. The output layer applies a softmax function to the splice site strength score from the scoring network. The softmax layer itself does not contain any variables, so it is compatible with any number of alternative splice sites.

### 3.3 Scoring Network

The scoring network takes inputs derived from one alternative splice site and outputs an unnormalized strength score. Let *s*_*i*_ denote the strength of the *k*-th alternative splice site such that 

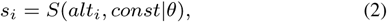

where *S* denotes the scoring network and *θ* represents the parameters of the scoring network *S*. Importantly, the parameters *θ* do not depend on *k*, but are shared between all splice sites.

We implement four different architectures for the scoring network and evaluate their performance carefully. The simpler architectures we evaluate use scoring networks that are independent, such that the competitive behavior of the model is achieved only through the requirement that the *PSI*_*k*_, *k ∈* 1, 2, …, *K* sum to 1.

However, we also evaluate a number of architectures in which we allow *lateral* connections between the alternative splice sites within the same event. In particular, we utilize *communication networks* (Sukhbaatar *et al*., 2016) and LSTMs, which are described in Sections 3.3.2, 3.3.3, and 3.3.4 respectively.

#### 3.3.1 Independent scoring networks

These types of networks take as input a set of splice site sequences and predict a scalar unnormalized score as in Eq (2) without any lateral connections to the other scoring networks. We use an architecture in which we use a stack of multiple convolutional layers on each of the input sequences. The outputs of each stack are concatenated with the intron length feature and followed by multiple fully connected layers.

Figure 5c shows the architecture of the independent convolutional scoring network. We use three columns of stacked convolutions with independent parameters that each use one of the three sequence windows described in Section 3.2 as input. A *convolution module* contains the convolutional layer itself, followed by a ReLU non-linearity, and a batch-normalization layer (Ioffe and Szegedy, 2015). We do not use pooling, because some filters, like the core splice site motif, can be highly sensitive to the precise location and are not invariant to shifts.

**Fig. 5.**
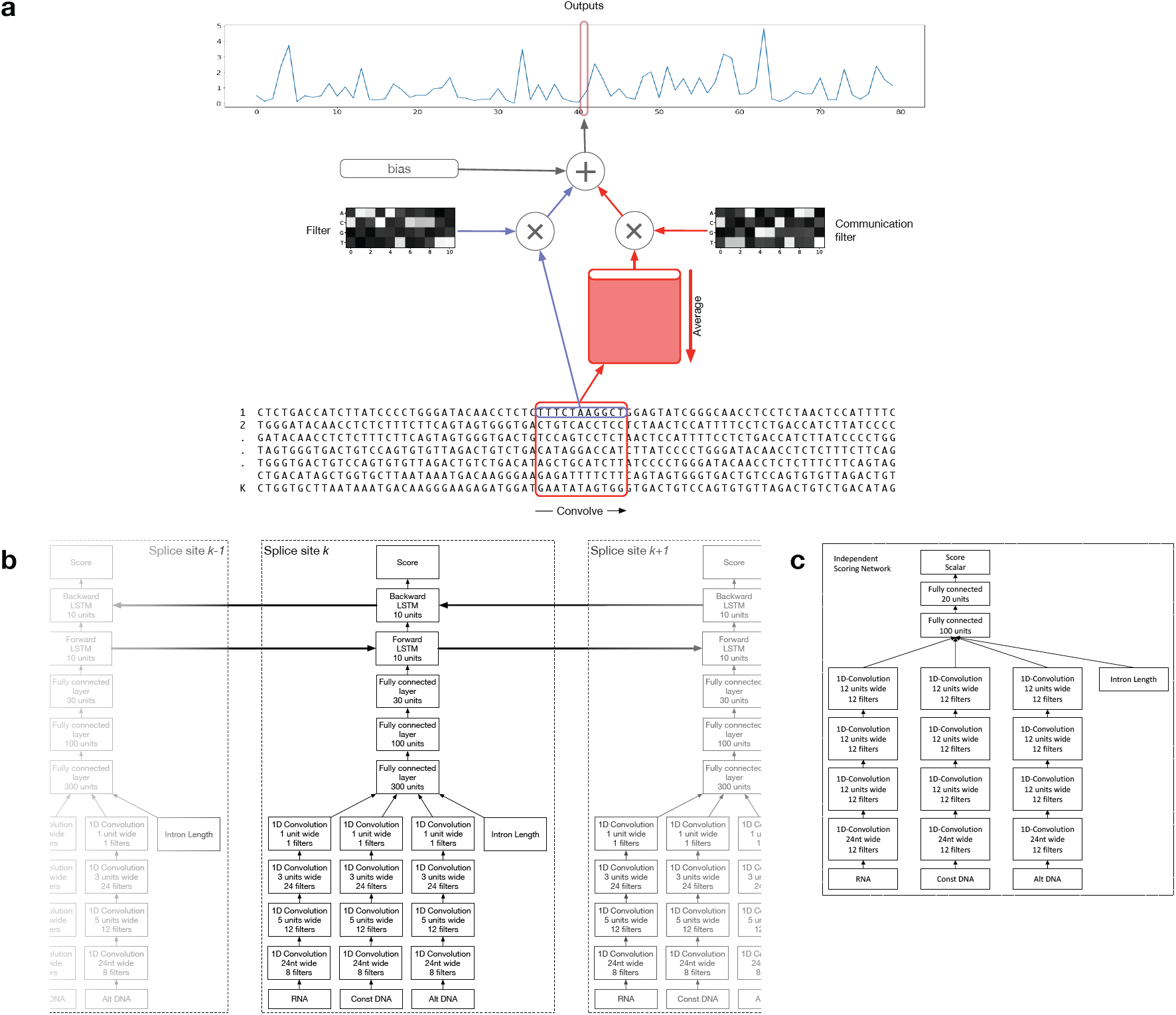
Model architectures. (a) Convolutional layer with communication: This is a variant of the typical 1-D convolutional layer in case we apply a convolution to multiple sequences such as a set of alternative splice site. For a given sequence *k* = 1, …, *K*, the output of this layer is the sum of a bias term, the filter responses to the sequence *k*, and the responses to a second set of filters applied to the average of all inputs across *k*. (b) Architecture of our bidirectional LSTM Model: Features for splice site *k* are extracted from the DNA/RNA sequences using a series of convolutional modules. The filtermaps for the different sequences are concatenated with the intron length feature. The concatenated features are then further processed by several fully connected modules and two LSTM cells. The forward cell is connected to splice site *k* − 1 and the backwards cell is connected to splice site *k* + 1. (c) Independent convolutional scoring network: Scoring networks consist of several convolutional layers followed by fully connected layers and are not laterally connected.

Following any number of convolution modules, the final filtermaps from the three convolution columns are flattened and concatenated with the intron length feature. These activations are the input to the following sequence of fully connected modules, each consisting of a fully connected layer, a ReLU function, and a batch-normalization layer.

The final fully connected module, as in all other architectures, has an output size of one to connect to the softmax output layer.

#### 3.3.2 Communication networks

When the scoring networks for competing splice sites are independent, they can only interact linearly through the softmax layer in the output network. We use an approach that is similar to Communication networks (Sukhbaatar *et al*., 2016) to model more complicated interactions between the splice sites, which we adapt to convolutional layers.

In our version of communication networks, a convolutional layer uses two sets of filters: one which operates on the current splice site and one which operates on a shared buffer that contains the average inputs of all splice sites. Each splice site’s consecutive hidden layer then takes both the shared communication buffer’s activations and the activations from the splice site’s own previous hidden layer as input.

We apply communication to the convolutional layers in the following way: the output of a convolution layer with communication for a given splice site *k* is the sum of a global bias term, the response of a filter to the *k*-th input and the response of a second set of filters to the inputs averaged across all splice sites 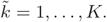

The output *y*_*k,g,j*_ for splice site *k*, output filter *g* at position *j* is

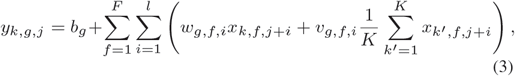

where *b*_*g*_ is the bias of the *g*-th filter, *l* is the filter width, *w*_*g,f,i*_ is the filter value of the *g*-th filter to the *f* -th channel at position *i, x*_*k,f,j*+*i*_ is the input value of splice site *k* in the *f*-th channel at the *j* + *i*-th position, *v_f;i_* is the weight of the *g*-th communication filter to the *f*-th channel, at position *i*, and 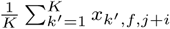*1* is the input averaged across all splice sites in channel *f* at the *j* + *i*-th position.

Figure 5a further illustrates the communication network concept.

#### 3.3.3 Output LSTM

Communication networks are one possibility for modeling the interaction between splice sites. However, an alternative that explicitly takes the ordering of the splice sites into account are *Long Short-Term Memory* (LSTM) networks (Hochreiter and Schmidhuber, 1997). LSTM networks are a type of recurrent neural network architecture, that uses memory cells with gates that control the flow of information and can be learned by backpropagation.

LSTMs can be used on any kind of sequential or time-series data such as strings of words in machine translation or frames of speech recordings. For the purpose of alternative splicing prediction we apply LSTMs to the sequence of splice sites.

We implement a hybrid architecture in which we keep the lower convolutional layers from the independent scoring networks (Section 3.3.1) and replace the model’s fully connected layers with a bidirectional LSTM which consists of one LSTM running from 3’ to 5’ and the other one from 5’ to 3.

Figure 5b shows the architecture we have chosen. As in the independent scoring network, three different sequence windows from each splice site are first processed by a series of 1D convolutional modules before being concatenated with the intron length and are further passed through several fully connected modules. However, the signal from splice site *k* is now further processed by two LSTM cells which will connect to the adjacent splice sites *k −* 1 and *k* + 1, allowing for propagation of information between them.

#### 3.3.4 Resnet + LSTM

Resnets (He *et al*., 2015) are a class of models that enable training neural networks that are much deeper than previously possible by explicitly reformulating the layers as learning residual functions with reference to the layer inputs. Rather than learning a mapping *H*(*x*) directly, aresnet implements the residual mapping *F* (*x*) = *H*(*x*) *− x*. The desired mapping *H*(*x*) can then be reformulated as *F* (*x*) + *x*. In practice, resnets add shortcut connections that skip one or several layers. These shortcut connections can be added almost anywhere; for example, around convolutional or fully connected layers.

In our implementation we replace the convolutional layers from the LSTM model (Section 3.3.3) with the Resnet-26 model (He *et al*., 2015), while keeping the higher level fully-connected layers and LSTM the same. Table 2 shows the parameters of the Resnet-26 architecture.

**Table 2.** Resnet-26 architecture.

| Bottleneck Block | Parameters |  |  |  |
| --- | --- | --- | --- | --- |
|  | width | out_dim | oper_dim | stride |
| 1 | 24 | 8 | 1 | 1 |
| 2 | 8 | 32 | 8 | 1 |
| 3 | 8 | 64 | 16 | 2 |
|  | 8 | 64 | 16 | 1 |
| 4 | 8 | 128 | 32 | 2 |
|  | 8 | 256 | 64 | 1 |
| 5 | 8 | 256 | 64 | 1 |
|  | 8 | 256 | 64 | 1 |

This residual network replaces the convolutional layers in the LSTM model for the Resnet+LSTM model. Each bottleneck block can be skipped with a residual connection. Within each bottleneck block there are one or more convolutional layers parametrized by *width* (the width of the convolutional filter), *out\_dim* (the number of output filters), *oper\_dim* (the number of filters in the middle of the bottleneck block), and *stride* (the stride of the convolution operation).

### 4 Results

### 4.1 Genome-wide Performance

We trained each of the four models presented above on both our acceptor and donor datasets. We use five-fold cross-validation by splitting our datasets according to the following method: using a transcript database, we start with a set of intervals that each span two consecutive genes initially. Then we iteratively merge all regions that are less than 250nt apart until merging is no longer possible. This results in a set of 175 distinct genomic regions. We randomly split the resulting genomic regions into five folds for cross validation. Each fold uses 135 regions for training, five regions for validation, and 35 regions as the test set. When training each fold, we use the validation set for early stopping and hyperparameter optimization. This procedure results in more balanced splits than splitting by entire chromosomes, which have very different lengths.

Figure 6 plots the cross-entropy error over time during training. It is evident that the LSTM and Resnet models are able to fit the training data significantly better and achieve a lower final training loss.

**Fig. 6.**
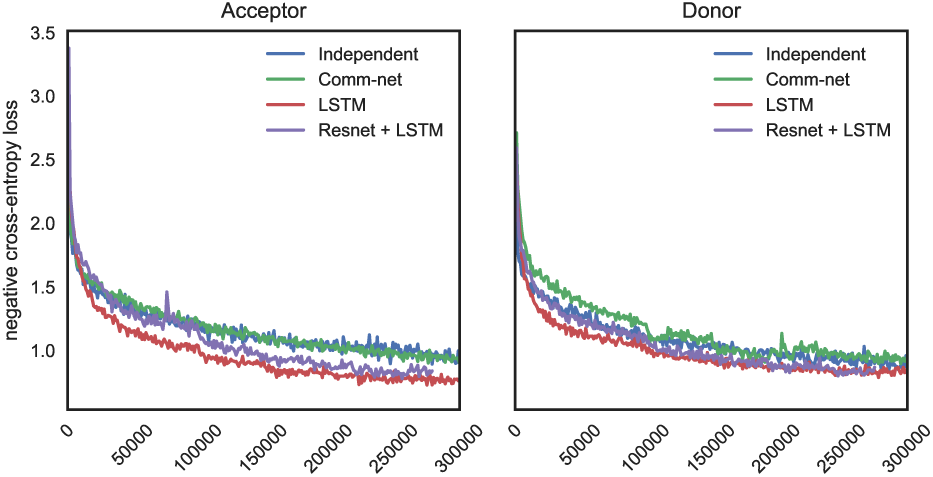
Training cross-entropy loss.

**Fig. 7.**
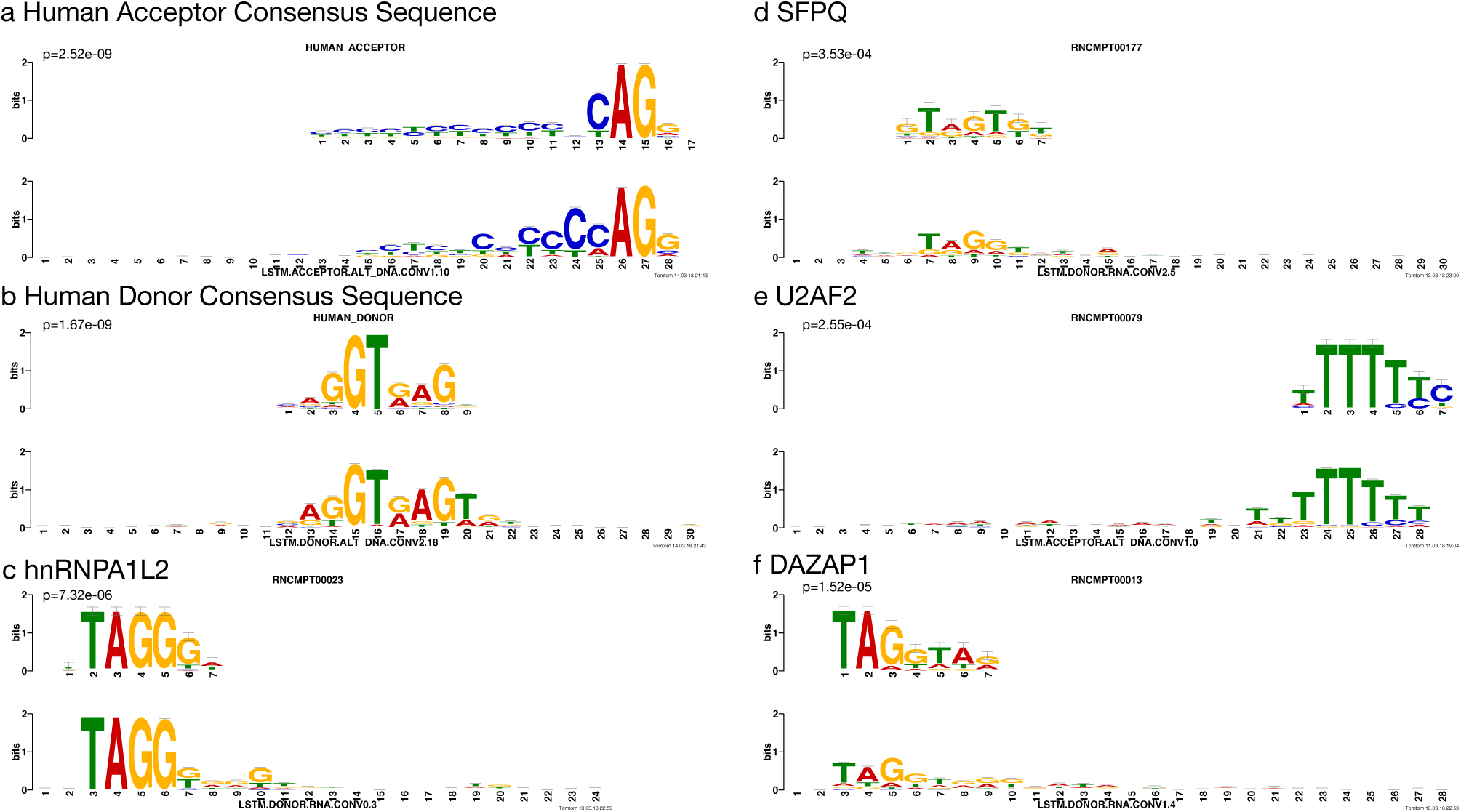
Motif visualization. These are matches of motifs learned by the LSTM acceptor and donor models against known splicing regulators. Motifs are extracted using the method by Alipanahi et al. (2015) and then referenced against the human splice site acceptor consensus motifs (Zhang, 1998) and RNA binding elements from RNA compete (Ray et al., 2013) with TOMTOM (Gupta et al., 2007). Reference motifs are on the top and matching motifs learned by COSSMO are on the bottom. The p-values are reported by the TOMTOM algorithm.

We compute the accuracy, loss and *R*^2^ on each fold and report the mean score across the folds as well as the standard error. Table 3 shows the accuracy of the different COSSMO models, as well as MaxEntScan, on the same data. We define the accuracy as the frequency with which COSSMO correctly predicts the splice site with the maximal PSI value.

**Table 3.** Accuracy of COSSMO models and MaxEntScan

| Model | All events |  | Alternatively spliced |  |
| --- | --- | --- | --- | --- |
| | Acceptor<br>$n = 172,940$ | Donor<br>$n = 175,000$ | Acceptor<br>$n = 22,564$ | Donor<br>$n = 25,865$ |
| MaxEntScan | 0.322 | 0.371 | 0.181 | 0.210 |
| Independent | 0.520 ($\pm 0.004$) | 0.569 ($\pm 0.003$) | 0.331 ($\pm 0.003$) | 0.376 ($\pm 0.002$) |
| Comm-net | 0.540 ($\pm 0.008$) | 0.653 ($\pm 0.001$) | 0.350 ($\pm 0.007$) | 0.482 ($\pm 0.005$) |
| LSTM | **0.700** ($\pm 0.004$) | **0.711** ($\pm 0.002$) | **0.533** ($\pm 0.006$) | **0.548** ($\pm 0.005$) |
| Resnet + LSTM | 0.698 ($\pm 0.005$) | 0.701 ($\pm 0.003$) | 0.526 ($\pm 0.008$) | 0.540 ($\pm 0.005$) |

Accuracy is defined as the frequency at which the model correctly identifies the splice site with maximum PSI. Accuracies are computed as the average of five cross-validation folds with standard errors given in parentheses. The right columns shows accuracy only for those events that contain two or more annotated splice sites.

It is not surprising that MaxEntScan, which can predict the strongest splice site with a probability of 32.2% (acceptor), or 37.1% (donor), is outperformed by even the simple independent COSSMO model (52.0% on acceptor and 56.9% on donor). After all, MaxEntScan uses a smaller sequence window, is trained on a smaller, different, dataset, and does not utilize the competitive training procedure.

Our best model, which uses an LSTM, achieves 70.0% on the acceptor dataset and 71.1% on the donor. Despite being a deeper, more powerful model, the resnet with an LSTM achieves slightly worse performance than the “LSTM only” model. Our interpretation of this result is that the LSTM layer is critical for good performance but the convolutional subnetwork in the LSTM network is sufficient to learn useful features. As a result, the deeper feature extraction network in the LSTM+Resnet does not achieve better performance than the slightly simpler LSTM model.

Accuracy is intuitive to interpret but it only takes into account whether or not COSSMO predicts the strongest splice site in a large set correctly. Accuracy does not take into account how well the predictions fit the PSI distribution overall. Tables 4 shows the cross-entropy error of the four different COSSMO models and Table 5 shows the coefficient of determination *R*^2^. While MaxEntScan performs relatively well at predicting the dominant splice, as shown by the accuracy, it performs much worse at predicting the PSI distribution in general. This should not be a surprise given that MaxEntScan was designed to score splice sites in isolation. Still, this demonstrates the need to model splice sites in their local context if predicting their relative utilization is the goal.

**Table 4.**
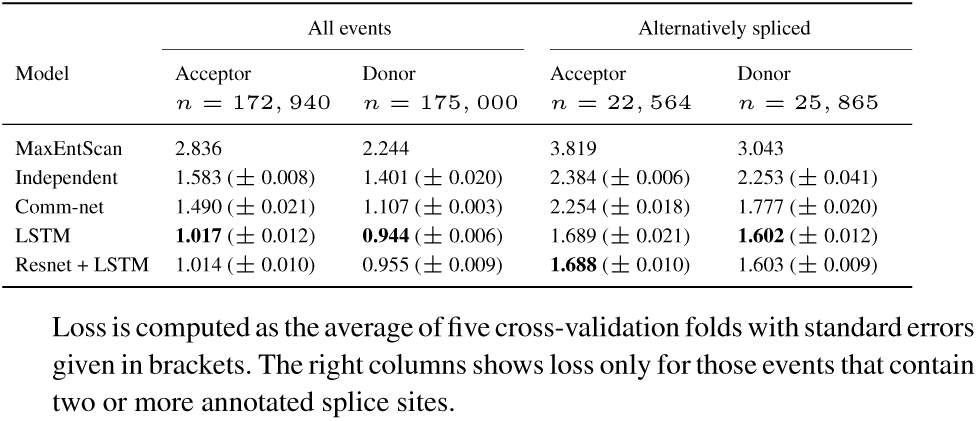
Cross-entropy error of COSSMO models and MaxEntScan.

**Table 5.** Coefficient of determination (*R*^2^) of COSSMO models and MaxEntScan

| Model | All events |  | Alternatively spliced |  |
| --- | --- | --- | --- | --- |
| | Acceptor<br>$n = 172,940$ | Donor<br>$n = 175,000$ | Acceptor<br>$n = 22,564$ | Donor<br>$n = 25,865$ |
| MaxEntScan | 0.080 | 0.195 | -0.105 | 0.038 |
| Independent | 0.389 ($\pm 0.003$) | 0.438 ($\pm 0.005$) | 0.187 ($\pm 0.003$) | 0.208 ($\pm 0.009$) |
| Comm-net | 0.421 ($\pm 0.007$) | 0.552 ($\pm 0.001$) | 0.229 ($\pm 0.003$) | 0.372 ($\pm 0.008$) |
| LSTM | **0.614** ($\pm 0.005$) | **0.628** ($\pm 0.003$) | **0.430** ($\pm 0.006$) | **0.453** ($\pm 0.005$) |
| Resnet + LSTM | 0.612 ($\pm 0.006$) | 0.618 ($\pm 0.004$) | 0.420 ($\pm 0.007$) | 0.448 ($\pm 0.004$) |

The coefficient of determination is computed using the class prior $1/K$ (uniform distribution over alternative splice sites) as baseline. For COSSMO, $R^2$ is computed as the average of five cross-validation folds with standard errors given in parentheses. The right columns shows loss only for those events that contain two or more annotated splice sites.

### 4.2 Performance on Alternatively Spliced Events

The majority of the events in our dataset have only one annotated splice site (Figure 2). To gain a deeper understanding of COSSMO’s performance, it also helps to stratify the dataset and only look at events that are alternatively spliced (have more than one *annotated* splice site). The performance of MaxEntScan and COSSMO on this subset is shown in the right hand columns of Tables 3, 4, and 5. This is a much more challenging subset of the data and performance of MaxEntScan and COSSMO is both impacted when more than one annotated site is present. However, while MaxEntScan accuracy drops by almost half on this subset, COSSMO’s relative drop in performance is lower, with the more complicated models (“LSTM” and “LSTM + Resnet”) seeing a smaller relative drop in accuracy than the simpler models (“Independent” and “Comm-net”).

Table 5 shows that MaxEntScan breaks down on the alternatively spliced subset. On the alternative acceptor dataset, MaxEntScan explains less of the variance than simply predicting a uniform distribution over splice sites, manifesting as a negative *R*^2^ score. The *R*^2^ of the COSSMO LSTM model drops from 61% to 43% on this more difficult subset.

### 4.3 Error analysis

It is helpful to examine the types of errors our predictors make and how the models differ in their mistakes. Figure 8 shows a breakdown of the types of splice site out of the different categories presented in Table 1. The bars show the proportions of the types of the predicted dominant splice site. As expected, the annotated splice sites almost always have greater true PSI than *de-novo* or decoy splice sites in our dataset.

**Fig. 8.**
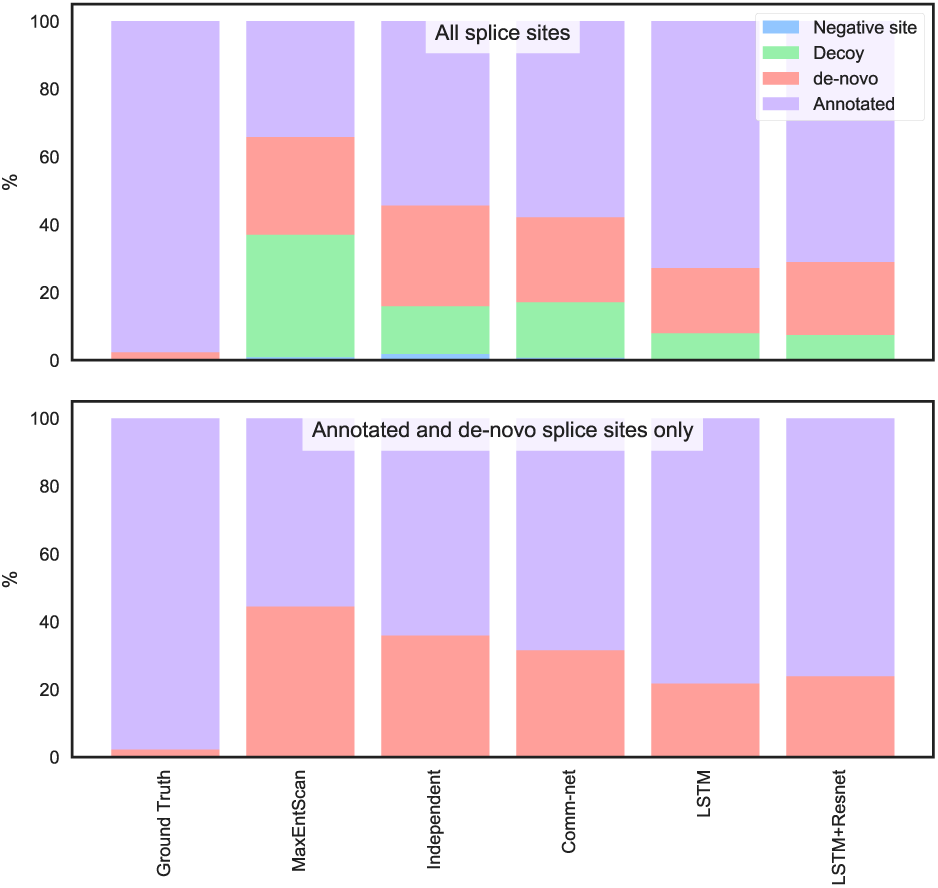
Type of splice site predicted by different models. This plot shows how often each model predicts a splice site of each possible type to be the strongest as opposed to the ground truth. The top plot uses real and non-splice sites as alternatives, while the bottom plot uses only real splice sites, which is less a challenging task for the model. Only the alternative acceptor data is shown.

Our best model, which use an LSTM almost never mistakes a negative site for real. However, it can be fooled by a decoy site approximately 8% of the time. The LSTM model also predicts a *de-novo* site to be dominant approximately 20% of the time. The simpler independent and communication network COSSMO models make the same errors more often.

MaxEntScan incorrectly predicts a decoy splice site to be dominant nearly 40% of the time. This is not surprising since the decoy sites are chosen to be cases that MaxEntScan scores higher than they should be according to our RNA-Seq data and the data is thus biased to producing wrong predictions from MaxEntScan. The results, however, show that our dataset design is relatively successful in producing models that are less likely to be fooled by cryptic splice sites and other sequences that are not recognized by the spliceosome, even though they look very similar to true splice sites.

To examine how much our dataset design disadvantages MaxEntScan, we remove both decoy and negative sites from the test set and use each model to predict the dominant site only from the annotated and *de-novo* sites. This is a much easier prediction problem because in this setting, the model must only choose between an average of five putative splice sites rather than around 80 sites in the full setting.

Even in this scenario, MaxEntScan incorrectly predicts a *de-novo* site to be dominant in about 44% of test cases, while the LSTM COSSMO model has about half the error rate, demonstrating that the performance gap persists even when we correct for possible bias in the dataset design.

## 5 Model interpretation

To visualize the motifs COSSMO learns we follow a similar procedure to Alipanahi *et al*. (2015). We run approximately 2M splice sites from our test set through the model and extract the activations after each convolutional layer, just after applying the ReLU function. We then threshold those activations at the 99.9% percentile to keep only those input sequences to which the filter responds the most strongly. Then, we align the input sequences according to the position of the output unit and compute the position probability matrix corresponding to each filter.

For both the acceptor and donor LSTM models, we extract motifs learned for the alternative DNA, constant DNA, and spliced RNA input sequences by the first, second, and third convolutional layer.

We then use TOMTOM (Gupta *et al*., 2007) to find matches of the motifs learned by COSSMO in the RNA compete database of RNA-binding elements (Ray *et al*., 2013) and against the human acceptor and donor site consensus motifs (Zhang, 1998). We find that COSSMO’s learned motifs match a large number of the most important known splicing factor binding sites as well as the acceptor and donor consensus motifs.

Figure 7 shows several examples of high-certainty matches of COSSMO’s motifs against known motifs. In particular, COSSMO learns motifs that almost perfectly match the known acceptor and donor site consensus motifs (a & b). TOMTOM produces a large number of matches against the RNAcompete motifs, among them many of the most important splicing factors (*e.g*. hnRNPA2B1, hnRNPA1L2, hnRNPA1, hnRNPH2, PTBP1, QK1, SFPQ, SRSF1, SRSF2, SRSF7, SRSF9, SRSF10, U2AF2, YBX1). In total we find 83 matches against RNAcompete motifs with *p <* 10^*−* 2^ in the acceptor model and 140 matches in the donor model. Figure 7 shows some examples of strong matches to known splicing factors:

- hnRNPA1L2 (c, *p* = 7.32*e −* 06), which is a member of the hnRNP family of RNA-protein complexes that are involved in splicing control (Martinez-Contreras *et al*., 2007),
- SFPQ (d, *p* = 3.52*e −* 04), an essential pre-mRNA splicing factor required early in spliceosome formation and for splicing catalytic step II (Patton *et al*., 1993),
- U2AF2 (e, *p* = 2.55*e −* 04), which is a necessary part of the spliceosome and binds to the polypyrimidine tract (Zamore *et al*., 1992), and
- DAZAP1 (f, *p* = 1.52*e −* 05), which can activate weak exons by neutralizing splicing inhibitors (Choudhury *et al*., 2014).

The full list of motif matches is available in the supplementary materials.

## 6 Discussion

In this work, we introduced COSSMO, a computational model that enables accurate prediction of competitive alternative splice site selection from sequence alone. We describe how we generate a genome-wide dataset for training and evaluation by combining positive splice site examples from genome annotations and large-scale RNA-Seq datasets, as well as negative examples from random genomic background sequences, and decoy splice sites that receive high scores from MaxEntScan but lack evidence of splice site usage from RNA-Seq data.

We design four neural network architectures that adapt to a variable number of alternative splice sites and carefully evaluate them using genome-wide cross-validation. We find that all of our models performed better than MaxEntScan, but we also find large performance differences between the different COSSMO architectures. Independent scoring networks achieve decent performance but the best performance depends on a communication mechanisms between the scoring networks. Of these models, the recurrent LSTM model achieved better accuracy than the communication network, which does not take the ordering of the splice sites into account.

We also demonstrate that COSSMO learns *ab-initio*, without any feature engineering or built-in knowledge, a wide array of binding motifs that correspond to the splice site consensus sequences and known splicing factors.

Our work demonstrates that it is possible to use deep learning to predict splice site choice with high accuracy, which can be extended to predict how genomic variation affects splice site choice through mechanisms like splice site variants or cryptic splice site activation.

## References

Alipanahi, B., Delong, A., Weirauch, M. T., and Frey, B. (2015). Predicting the sequence specificities of DNA-and RNA-binding proteins by deep learning. Nature Biotechnology, 33(8), 831–838.

Barash, Y., Calarco, J. A., Gao, W., Pan, Q., Wang, X., Shai, O., Blencowe, B. J., and Frey, B. J. (2010). Deciphering the splicing code. Nature, 465(7294), 53–59.

Busch, A. and Hertel, K. J. (2015). Splicing predictions reliably classify different types of alternative splicing. RNA, 21(5), 813–823.

Choudhury, R., Roy, S. G., Tsai, Y. S., Tripathy, A., Graves, L. M., and Wang, Z. (2014). The splicing activator dazap1 integrates splicing control into mek/erk-regulated cell proliferation and migration. Nature Communications, 5, 3078 EP–.

GTEx Consortium, T. (2013). The Genotype-Tissue Expression (GTEx) project. Nature Genetics, 45(6), 580–585.

Gupta, S., Stamatoyannopoulos, J. A., Bailey, T. L., and Noble, W. S. (2007). Quantifying similarity between motifs. Genome Biology, 8(2), R24.

Harrow, J., Frankish, A., Gonzalez, J. M., Tapanari, E., Diekhans, M., Kokocinski, F., Aken, B. L., Barrell, D., Zadissa, A., Searle, S., Barnes, I., Bignell, A., Boychenko, V., Hunt, T., Kay, M., Mukherjee, G., Rajan, J., Despacio-Reyes, G., Saunders, G., Steward, C., Harte, R., Lin, M., Howald, C., Tanzer, A., Derrien, T., Chrast, J., Walters, N., Balasubramanian, S., Pei, B., Tress, M., Rodriguez, J. M., Ezkurdia, I., van Baren, J., Brent, M., Haussler, D., Kellis, M., Valencia, A., Reymond, A., Gerstein, M., Guigó, R., and Hubbard, T. J. (2012). Gencode: The reference human genome annotation for the encode project. Genome Research, 22(9), 1760–1774.

He, K., Zhang, X., Ren, S., and Sun, J. (2015). Deep residual learning for image recognition. abs/1512.03385.

Hochreiter, S. and Schmidhuber, J. (1997). Long short-term memory. Neural Computation, 9(8), 1735–1780.

Hong, X., Scofield, D. G., and Lynch, M. (2006). Intron size, abundance, and distribution within untranslated regions of genes. Molecular Biology and Evolution, 23(12), 2392–2404.

Ioffe, S. and Szegedy, C. (2015). Batch normalization: Accelerating deep network training by reducing internal covariate shift. arXiv, abs/1502.03167.

Kelley, D. R., Reshef, Y. A., Belanger, D., McLean, C., Snoek, J., and Bileschi, M. (2018). Sequential regulatory activity prediction across chromosomes with convolutional neural networks. bioRxiv.

Kim, D., Langmead, B., and Salzberg, S. L. (2015). HISAT: a fast spliced aligner with low memory requirements. Nature Methods, 12(4), 357–360.

Leung, M. K. K., Xiong, H. Y., Lee, L. J., and Frey, B. J. (2014). Deep learning of the tissue-regulated splicing code. Bioinformatics (Oxford, England), 30(12), i21–i29.

Martinez-Contreras, R., Cloutier, P., Shkreta, L., Fisette, J.-F., Revil, T., and Chabot, B. (2007). hnrnp proteins and splicing control. Adv Exp Med Biol, 623, 123–147.

Patton, J. G., Porro, E. B., Galceran, J., Tempst, P., and Nadal-Ginard, B. (1993). Cloning and characterization of psf, a novel pre-mrna splicing factor. Genes Dev, 7(3), 393–406.

Quang, D. and Xie, X. (2016). DanQ: a hybrid convolutional and recurrent deep neural network for quantifying the function of DNA sequences. Nucleic Acids Research, 44(11), e107.

Ray, D., Kazan, H., Cook, K. B., Weirauch, M. T., Najafabadi, H. S., Li, X., Gueroussov, S., Albu, M., Zheng, H., Yang, A., Na, H., Irimia, M., Matzat, L. H., Dale, R. K., Smith, S. A., Yarosh, C. A., Kelly, S. M., Nabet, B., Mecenas, D., Li, W., Laishram, R. S., Qiao, M., Lipshitz, H. D., Piano, F., Corbett, A. H., Carstens, R. P., Frey, B. J., Anderson, R. A., Lynch, K. W., Penalva, L. O. F., Lei, E. P., Fraser, A. G., Blencowe, B. J., Morris, Q. D., and Hughes, T. R. (2013). A compendium of RNA-binding motifs for decoding gene regulation. Nature, 499(7457), 172–177.

Scotti, M. M. and Swanson, M. S. (2015). Rna mis-splicing in disease. Nature Reviews Genetics, 17, 19 EP –.

Sukhbaatar, S., Szlam, A., and Fergus, R. (2016). Learning multiagent communication with backpropagation. In D. D. Lee, M. Sugiyama, U. V. Luxburg, I. Guyon, and R. Garnett, editors, Advances in Neural Information Processing Systems 29, pages 2244–2252. Curran Associates, Inc.

Turunen, J. J., Niemelä, E. H., Verma, B., and Frilander, M. J. (2013). The significant other: splicing by the minor spliceosome. Wiley Interdisciplinary Reviews: RNA, 4(1), 61–76.

Vaz-Drago, R., Custódio, N., and Carmo-Fonseca, M. (2017). Deep intronic mutations and human disease. Human Genetics, 136(9), 1093–1111.

Wang, Z. and Burge, C. B. (2008). Splicing regulation: from a parts list of regulatory elements to an integrated splicing code. RNA, 14(5), 802–813.

Xiong, H. Y., Barash, Y., and Frey, B. J. (2011). Bayesian prediction of tissue-regulated splicing using RNA sequence and cellular context. Bioinformatics (Oxford, England), 27(18), 2554–2562.

Xiong, H. Y., Alipanahi, B., Lee, L. J., Bretschneider, H., Merico, D., Yuen, R. K. C., Hua, Y., Gueroussov, S., Najafabadi, H. S., Hughes, T. R., Morris, Q., Barash, Y., Krainer, A. R., Jojic, N., Scherer, S. W., Blencowe, B. J., and Frey, B. (2015). The human splicing code reveals new insights into the genetic determinants of disease. American Association for the Advancement of Science. Science.

Xiong, H. Y., Lee, L. J., Bretschneider, H., Gao, J., Jojic, N., and Frey, B. J. (2016). Probabilistic estimation of short sequence expression using rna-seq data and the positional bootstrap. bioRxiv.

Yeo, G. and Burge, C. B. (2004). Maximum entropy modeling of short sequence motifs with applications to RNA splicing signals. Journal of Computational Biology, 11(2–3), 377–394.

Zamore, P. D., Patton, J. G., and Green, M. R. (1992). Cloning and domain structure of the mammalian splicing factor u2af. Nature, 355(6361), 609–614.

Zhang, M. Q. (1998). Statistical features of human exons and their flanking regions. Human Molecular Genetics, 7(5), 919–932.

